# Decellularized Banana Leaves: Eco-Friendly Scaffolds for Cell-Based Seafood

**DOI:** 10.1101/2023.11.03.565310

**Authors:** Reyhaneh Sarkarat, Amiti Banavar, Arian Amirvaresi, Xinxin Li, Cuong Nguyen, David L. Kaplan, Nitin Nitin, Reza Ovissipour

## Abstract

Cellular agriculture, as an emerging food production system, holds potential to address sustainability, food security, and agricultural resilience. Within the cell-based meat supply chain, one of the key steps is scaffolding. In this study, we assessed decellularized banana leaves, various coating materials, and different cell seeding strategies to determine their effects on cell viability, cell growth, cell alignment, and the response of the materials to thermal processing. The efficiency of decellularization was verified through DNA quantification, which decreased from 445 ng/mg in fresh banana leaves to non-detectable levels in the decellularized samples. This was further confirmed by FTIR and PCA modeling. Cell viability exceeded 98% on uncoated, soy-coated, and gelatin-coated samples of the decellularized banana leaves. Alignment of cells on gelatin-coated samples was the highest among the samples, with a dominant orientation of 65.8°, compared to soy-coated and uncoated samples with dominant orientations of 9.2° and −6.3°, respectively. In terms of quality attributes, the kinetics of shrinkage indicated that coating with soy and the presence of cells increased the activation energy due to the higher energy required for protein denaturation. Moreover, the kinetics of area changes in plain scaffolds without cells followed a first-order pattern, while with seeded cells a second-order pattern was followed. In summary, decellularized banana leaves present a sustainable and suitable biomaterial to support cells towards future needs related to meat production.

## Introduction

The continuous growth of the global population places significant demands on traditional farming practices. Projections indicate that the world’s population is expected to reach between 9 and 11 billion people by the year 2050 from the current level of eight billion. This demographic expansion amplifies pressure on existing agricultural systems (Eibl et al., 2021; Röös et al., 2017). This population growth necessitates the provision of food and agricultural commodities, which must be accomplished within the constraints of limited arable land and the looming challenges posed by climate change. In order to attain this objective, it is imperative to make a significant advancements in agricultural production within the foreseeable future (Rischer et al., 2020). One potential option or alternative involves the adoption of cellular agriculture, which presents a more sustainable and ecologically conscious approach to agricultural production.

Cellular agriculture, also known as cultured or lab-grown agriculture, is a cutting-edge field that aims to produce animal-derived food products without traditional farming practices. The field involves using cell culture and tissue engineering techniques to grow and multiply animal cells in a controlled laboratory or factory setting, ultimately generating proteins and muscle and fat tissues for meat, fish, dairy, and egg products (Soice & Johnston, 2021).

Cellular agriculture offers numerous advantages for sustainable and ethical food production. The approaches that are utilized reduce environmental impact by requiring fewer resources and emitting fewer greenhouse gases (Rubio et al., 2019). Animal welfare is also improved as the methods eliminate the need for raising and slaughtering animals. Furthermore, food safety is enhanced through the use of highly controlled production conditions, with the goal of reducing the risk of foodborne illnesses and the use of antibiotics or pesticides. Customization options also allow for control over nutritional content and flavor. Additionally, cellular agriculture can help conserve land and protect biodiversity (Kumar et al., 2021).

A fundamental feature in cultivated meat production is the process of biomaterials used for scaffolding. This feature involves seeding cells onto or into a biomaterial tissue-like platform known as a scaffold, which plays a crucial role in supporting the growth, differentiation, and proliferation of the cells (Bomkamp et al., 2022). There are two general approaches for scaffolding in tissue engineering, bottom-up and top-down. In the first approach, the scaffold is fabricated from the bottom-up using natural biomaterials or synthetic polymers, and the top-down approach uses modifications of prefabricated structures such as porous plant tissues to generate the scaffolds (Nichol & Khademhosseini, 2009). In order to use plant tissues, the extracellular matrix (ECM) should be isolated from the cellular materials without excessive damage to the structure and biochemical characteristics. Decellularized tissue from common plants can be considered a potential option for cultivated meat scaffolding, since these are edible, safe to consume, and biocompatible, while also offering important attributes such as texture and porosity (Jones et al., 2023).

Using chemical solutions such as acids and bases or ionic solutions and detergents are efficient strategies for decellularization. Choosing a chemical that successfully removes cells from the tissue with minimal damage to the ECM is a crucial step in the decellularization process. Since the final products will be edible, it is essential for the solutions used to be generally recognized as safe (GRAS) and regulated for use in the food industry. Sodium dodecyl sulfate (SDS) and Polysorbate-20 that are used in the present study are examples of detergents that are regulated by the US Food and Drug administration (FDA) and are employed in different steps in food processing (Jones et al., 2023).

In previous studies, the efficiency of using decellularized spinach leaves as scaffolds for growing bovine satellite cells and producing cultured meat was assessed (Jones et al., 2021). This scaffold structure supported some mammalian cell types with bovine satellite cells viable on the surface of decellularized spinach leaves for extended growth periods, while incorporating regulated detergents met food safety requirements (Jones et al., 2021). Cheng et al. (2020) assessed a range of decellularized fruit and vegetables for scaffolding, such as carrot, celery, broccoli, leek, cucumber, potato, apple, asparagus, and green onion. The results of quantitative analysis of C2C12 and human skeletal muscle cells alignment and differentiation on these structures revealed that decellularized green onion scaffold was a cost-effective substrate for cultivated meat production (Cheng et al., 2020). In another study, Perreault et al. (2023) investigated agricultural byproducts, such as corn husks and jackfruit rinds for scaffolds via decellularization, and suggested using these wastes as a sustainable substrate sources (Perreault et al., 2023). Aside from raw material sources for these decellularized scaffolds, consideration for cell seeding and food product development, including changes associated with cooking should also be considered (Ovissipour et al., 2017).

The primary objective of the present study was to assess the suitability of decellularized banana leaves as scaffolds for cultivated meat production. Various seeding techniques and coating materials were explored to optimize the process. Additionally, the quality of the materials was assessed using standard protocols after subjecting the products to pasteurization time and temperature schedules.

## Materials and Methods

Decellularization process Fresh banana leaves (*Musa* spp.) were obtained from the local grocery store, cleaned to remove dirt and debris, and disinfected using 70% ethanol across the surface of each sample. Banana leaves were cut into small pieces using a 8 mm biopsy punch. Samples were placed in a 100 mL beaker with 40 mL of 1% sodium dodecyl sulfate (SDS) (Thermo Scientific) solution and agitated in an orbital shaker at 100 RPM at ambient temperature for 3 days. This was followed by 3 washing steps with deionized water, allowing 5 minutes of shaking between washes. A solution of 10% bleach (Chlorox, 7.5% Sodium hypochlorite) and 1% Tween 20 (Thermo Fisher) was used as a detergent to remove the cells from the plant materials. The samples were then returned to the orbital shaker and agitated for 24-48 h, until the samples were colorless, as an indicator of complete cell removal. Samples were then washed with deionized water three times to remove any remaining bleach solution and dried for 10 minutes in a laminar flow hood. Decellularized samples freeze dried and stored at −80°C before characterization and use.

### Scaffold preparation

Decellularized scaffolds were removed from −80°C and allowed to thaw for 30 minutes at room temperature in a laminar flow hood. The scaffolds were then placed in 70% ethanol solution for sterilization for one h and then washed three times in PBS. Scaffolds in each coating group (0.1% porcine gelatine, and 0.1% soy protein isolate) were suspended in the coating solution for an additional 2 hours, while the uncoated (control) scaffolds were suspended in growth media.

### Cell culture

Zebrafish fibroblast stem cells (ZEM2S, CRL-2147™) were acquired from the American Type Cell Culture (ATCC), thawed, and subcultured to passage 5-10 before use in scaffold seeding. The thawing method consisted of allowing the ampoules to reach 28°C, at which point the cells were resuspended in growth media and centrifuged at 500 *g* for 7 minutes. The supernatant was removed and the cells were immediately resuspended in growth media. Cells were seeded at 4,000 cells/cm^2^ in T-75 flasks. The subcultures were initiated when each flask reached 70-85% confluency, and consisted of the removal of spent cell culture media, a single wash with phosphate buffered saline (PBS), and cell detachment via trypsinization. Trypsin-EDTA (1 mL, 0.025%) (Thermo Fisher) was distributed onto the cells and incubated at 28°C for 3-5 minutes until cells visibly detached. Trypsinized cells were transferred to a 15 mL falcon tube with 6 mL additional media, and subsequently centrifuged at 500 *g* for 7 minutes. The supernatant was removed and the remaining white pellet was resuspended in growth media. At this point, cells were counted using the Countess Automated Cell Counter (Thermo Fisher) and were then used in seeding experiments or seeded in T-75 flasks for further passage. For counting the cells, a 10 *μ*L cell suspension was combined with 10 *μ*L of Trypan blue dye and loaded onto each side of a cell counting chamber slide. Both live counts were taken and averaged to maximize consistency and accuracy for cell seeding. To prepare growth media, Leibovitz’s L15 (Gibco™, Grand Island, NY, USA) with phenol red indicator was first dissolved in 985 mL of sterile distilled water, and the pH of the media was adjusted to 7.8 via addition of 1 M HCl solution. The L15 media was filtered using a 0.22 *μ*m polyether sulfone (PES) filter (Millipore Sigma-Aldrich). The growth media consisted of 50 mL aliquots prepared every 2 weeks of 89% Leibovitz L-15 Medium, 9% Fetal Bovine Serum, and 2% anti-anti (antibiotic).

### Cell seeding

Seeding density was maintained at 300,000-500,000 cells suspended in 15-20 *μ*L of FBS per each scaffold in both natural and vacuum seeded groups. In the natural seeding process, each scaffold was seeded by directly pipetting the seeding solution onto each scaffold and allowing an incubation phase of 30 minutes for the attachment of the cells before gently adding 1 mL of growth media to each well. For the vacuum seeding process, the scaffolds were placed in 1.5 mL Eppendorf tubes before adding the seeding solution directly onto the surface of each scaffold. The tubes were briefly centrifuged at low speed to ensure that the scaffolds were in full contact with the seeding solution and no large bubbles were visible in the solution. The Eppendorf tubes were placed with the cap open in a vacuum bag, and the opening of each bag was folded over at least twice to ensure sterility while transferring tubes to the vacuum sealer. Each bag was vacuum sealed at 95% vacuum with a 5 second hold time, and then removed from the vacuum in sterile conditions. The scaffolds were then gently placed in each well of a 12 well plate with 1 mL of complete growth media in each well, and characterized immediately after seeding (0h), and after 24 to 48 h incubation at 28°C.

### Imaging

#### Scanning Electron Microscopy

Lyophilized scaffolds were sputter-coated with a 10 nm layer of Pt/Pd on a Leica ACE600 Sputter (Deerfield, IL, USA). Scaffolds were then imaged at an accelerating voltage of 5 kV on a FEI Quanta 600 FEG scanning electron microscope (Thermo Fisher). Pore size was measured using using ImageJ software Version 1.47a (National Institutes of Health (NIH), Rockville, MD, USA).

#### F-Actin cytoskeleton staining

ActinRed 555, and ActinGreen 488 (ReadyProbe) were applied separately. During pilot studies, one drop was added to each 2 mL well. The scaffolds were protected from light and incubated at room temperature for 30 minutes. The stain-containing solution was removed from the well, and the scaffolds were rinsed in PBS three times. DAPI (4′,6-diamidino-2-phenylindole) was used as a blue fluorescent nuclear stain. DAPI was diluted to 1 ng/mL and added to each well. The plates were incubated for 15 minutes, and DAPI containing solution was removed. The scaffolds were washed in PBS 3 times before being transferred to a glass slide for viewing.

#### Immunofluorescent staining

After fixation, samples were blocked with 3% Bovine serum albumin (Millipore Sigma, USA) for 30 min. The samples were incubated overnight with Paired Box 7 (PAX 7) antibodies (Invitrogen, USA) in blocking solution at 4°C. Afterwards, the samples were incubated in DAPI (1:1000, Thermo Fisher), AlexaFluor TM 488 phalloidin (1:400, Invitrogen) and Goat anti rabbit AlexaFluor 594 (1:400, Invitrogen) for 1 h at room temperature. Imaging by confocal laser scanning microscopy (CLSM) with 3D z-stack reconstruction was performed on the TCS SP8 microscope (Leica, Wetzlar, Germany).

#### Brightfield and confocal microscopy

An Olympus Inverted Microscope CKX53 with a phase contrast attachment was used to monitor cell growth and attachment. Fluorescence imaging was performed using a fluorescence imaging attachment at respective wavelengths for each stain. Before fluorescent staining, scaffolds were removed from the 12 well plates and placed into new plates to avoid imaging of cells attached to the wells versus the scaffolds. At each subsequent step, gentle pipetting was performed to the prevent detachment of cells. Scaffolds were washed with approximately 500 *μ*L of PBS per well. To fix the scaffolds, in 4% paraformaldehyde (PFA) solution in PBS was added to each well and scaffolds were incubated for 30 minutes at room temperature before removal of the PFA and resuspension in 500 *μ*L PBS.

### DNA Quantification

The Quant-iTTM PicogreenVR (PG) dsDNA Kit (Invitrogen, Orlando, USA) was used to determine the effectiveness of the decellularization process and the DNA content of the scaffolds at each time point, adapted from Dikici et al. (2019). Fresh banana leaf samples were cut using a 5 mm biopsy punch and 1 mm biopsy punch in the center of each sample. All other samples were decellularized and vacuum seeded as described above, then removed from growth media at the time of characterization. Samples were washed once with PBS and transferred to microtubes, snap frozen in liquid nitrogen, and stored at – 80°C for later quantification. Immediately upon removal from the −80°C freezer, 100 *μ*L of cell digestion buffer was added to each microtube. Cell digestion buffer consisted of 10 mM Tris-HCl, 1 mM ZnCl_2_, and 1% Triton-X100 in distilled water (dH_2_O). Each scaffold was ground into a fibrous pulp using microtube pestles until homogenous (minimal fiber visibly remaining), about 30 seconds per sample, and vortexed for 60 seconds, then kept overnight at 4°C. Picogreen (PG) working solution was prepared by diluting 20 Tris-EDTA (TE) 1:20 in dH_2_O (1 TE), then diluting the PG reagent 1:200 in 1 TE. Samples were subjected to 3 freeze-thaw (FT) cycles (10 min at −80°C and 20 min at 37°C, with 15 seconds vortexing between each cycle). The microtubes were centrifuged at 11,000 *g* for 5 minutes to separate fiber from lysate. Then, 100 *μ*L of the lysate was removed and combined thoroughly via pipetting with 100 mL PG working solution in Qubit tubes. Samples were then wrapped with aluminum foil and incubated at room temperature for 10 minutes. Fluorescence reading was conducted with the Qubit™ 4 Fluorometer (Thermo Fisher) at an excitation wavelength of 485 nm and an emission wavelength of 528 nm. The efficiency of decellularization was determined by comparing the DNA content (%) in fresh and decellularized leaves.

### Fourier-transform Infrared-Red

The spectra of three samples (n=3) of both fresh and decellularized banana leaves were collected using a Fourier-transform infrared spectrometer (FTIR) (PerkinElmer Universal ATR, PerkinElmer, Waltham, MA, USA). The spectrometer was coupled with a sampling accessory that included a Frontier Universal Diamond/ZnSe ATR crystal. The spectra were collected from 4000 cm^-1^ to 600 cm^-1^ with a 32-scan rate and a resolution of 4 cm^-1^. The samples were analyzed in triplicates, and means of the spectra was used for further analysis. Principal Component Analysis (PCA) was used via a loading plot constructed for PC1, which accounted for over 85% of the variables. This plot aimed to identify the most crucial variable responsible for the distinctions between the fresh (control) and decellularized banana leaves.

### Cell seeding efficiency

The efficiency of cell seeding was determined via DNA quantification and adapted from Buizler et al. (2013):

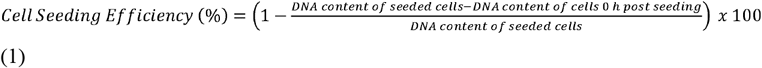

### Cell viability

Cell viability was measured after 48 h of cell seeding using a Live/Dead® staining kit (Thermo Fisher). The viability percentage was calculated using FIJI-Image 1.8.0_322 image process program (National Institutes of Health (NIH), Rockville, MD, USA), downloaded from https://imagej.net/Fiji, according to previous studies (Kothari et al., 2009; Schindelin et al., 2012).

### Cell alignment

Cell alignment was measured 48 h after cell seeding on the different scaffolds. After staining with DAPI, Actin and PAX7, the images were obtained using confocal laser microscopy as described above. The merged images were utilized for cell alignment measurements using the FIJI-Image 1.8.0_322 image process program by using an ellipse to each cell nucleus and cytoskeleton and measuring the angle of the maximum diameter relative to the horizontal axis of the image. The OrientationJ Plug-in was downloaded from www.epfl.ch/demo/orientation/ and added to the software to measure the orientation (Rezakhaniha et al., 2012; Puspoki et al., 2016; Jones et al., 2021). Color survey and 3D surface survey were created as well to visualize the orientation of cell cytoskeleton and the surface topography properties of each cell-seeded scaffold.

### Thermal treatment

Different sets of scaffolds, including uncoated and soy-coated vacuum-seeded samples were utilized to assess the kinetics of area shrinkage after cooking at different pasteurization temperatures (60, 65, 70, and 75°C) for varying durations, in accordance with the methods outlined previously in Ovissipour et al. (2017) (N=4). These times and temperatures were selected based on target bacteria and industry requirements to provide equivalent lethality and sufficient heating to inactivate *Listeria monocytogenes* (Ovissipour et al., 2017). Each scaffold was carefully positioned within an individually custom-built cylindrical aluminum test cell. These test cells were originally developed for kinetic studies of muscle foods (Ovissipour et al., 2017). Subsequently, the test cells were immersed in a water bath at each designated temperature point and retrieved at the specified time intervals, as detailed in Table 1. Following removal from the water bath, the aluminum sample cells were immediately submerged in an ice-water mixture to rapidly cool the samples and arrest the cooking process.

**Table 1:** Time and temperature schedule for thermal processing of scaffolds with and without cells.

| Temperature (°C) | Heating time (min) |  |  |  |  |  |
| --- | --- | --- | --- | --- | --- | --- |
| 60 | 1 | 2 | 4 | 8 | 10 | 12 |
| 65 | 1 | 2 | 3 | 4 | 5 | 6 |
| 70 | 0.5 | 1 | 2 | 3 | 4 | 5 |
| 75 | 0.25 | 0.5 | 0.75 | 1 | 1.5 | 2 |

#### Area Shrinkage

Area shrinkage was calculated based on the area of the individual sample before and after thermal processing, using ImageJ software Version 1.47a (National Institutes of Health (NIH), Rockville, MD, USA). The shrinkage ratio was calculated as:

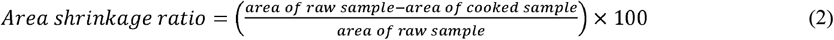

#### Kinetics Analysis

Quality degradation was calculated based on the area shrinkage. Reaction rates for quality degradation (area shrinkage) (*C*) in isothermal conditions are expressed in the following equation (Kong et al. 2007a):

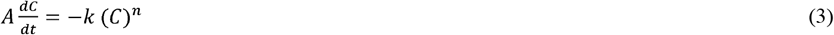

where *k* is the rate constant, *C* is the quality parameter (area shrinkage) at time *t*, and n is the order of reaction. To find the best empirical relationship, data was analyzed using zero-, first- and second-order kinetic models in as Eq. (4) – (6):zero-order:

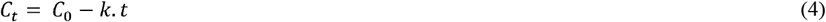

first-order:

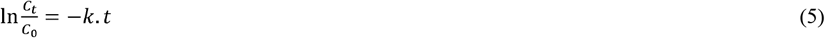

second-order:

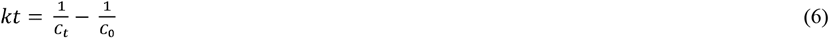

where *C*_*0*_ is the initial amount of the quality at time zero, *C*_*t*_ is the value at time *t, k* is the rate constant. In addition, Arrhenius equation was used to determine the degradation rate constant (*k*) on temperature, which is described as follows:

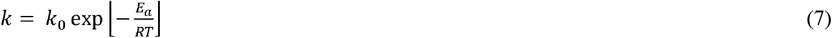

where *E*_*a*_ is the activation energy of the reaction (kJ/mol), *R* is universal gas constant (8.3145 J/mol/K), *T* is the absolute temperature (*K*) and *k*_*0*_ is the frequency factor (per min). By applying the Eq. (7) to a reaction, a plot of the rate constant on semi-logarithmic scale as a function of reciprocal absolute temperature (1/*T*) should yield a straight line. The activation energy can be determined as the slope of the line multiplied by the gas constant *R*.

### Statistical analysis

The data are presented as means for each treatment along with their respective standard deviations. A one-way ANOVA was conducted to assess the cell viability, cell adhesion and DNA content. To pinpoint any statistically significant distinctions among the treatment means, Tukey’s multiple comparison test was employed.

## Results and discussion

### Decellularization, FTIR spectra and DNA quantification

Banana leaves lost their color within 3 days after exposure to the decellularization solutions, and became translucent after 4 days (Figure 1A). To verify the efficiency of the decellularization process, DNA analysis was employed and indicated that the decellularization protocol reduced DNA from 445 ng/mg in the fresh banana leaves, to non-detectable level in the decellularized samples (Figure 1B) (n=9). The average pore size for the decellularized banana leaves was 147 ± 31 *μ*m (n=30 pores) (Figure 1C) which potentially provides a suitable platform for cell seeding and cell attachment and was in the range of pore sizes in decellularized plant tissues previously studied, which ranging from 50 to 300 *μ*m (Fontana et al., 2017; Jones et al., 2021; Thyden et al., 2022). While less than 50 ng/mg DNA is deemed sufficient for plant tissue decellularization for developing tissue in biomedical field (Crapo et al., 2011), for large-scale cultivated meat production and to reduce the environmental footprint and minimize chemical applications, adhering to this standard may not be required. This threshold was originally established to mitigate adverse host reactions to xenogeneic DNA from implanted decellularized tissue in patients. The efficiency of decellularization can vary depending on the type of plant tissue and the specific decellularization process employed. For instance, decellularizing spinach leaves required a 9-day process (Jones et al., 2021), while in the case of florets, only 48 hours were needed for complete decellularization (Thyden et al., 2022). In the present study, we were able to achieve complete decellularization in 4 days by implementing process modifications. Specifically, by introducing the small hole in the center of the banana leaves the duration of the decellularization process was reduced from the original 8 days to 4-days.

**Figure 1.**
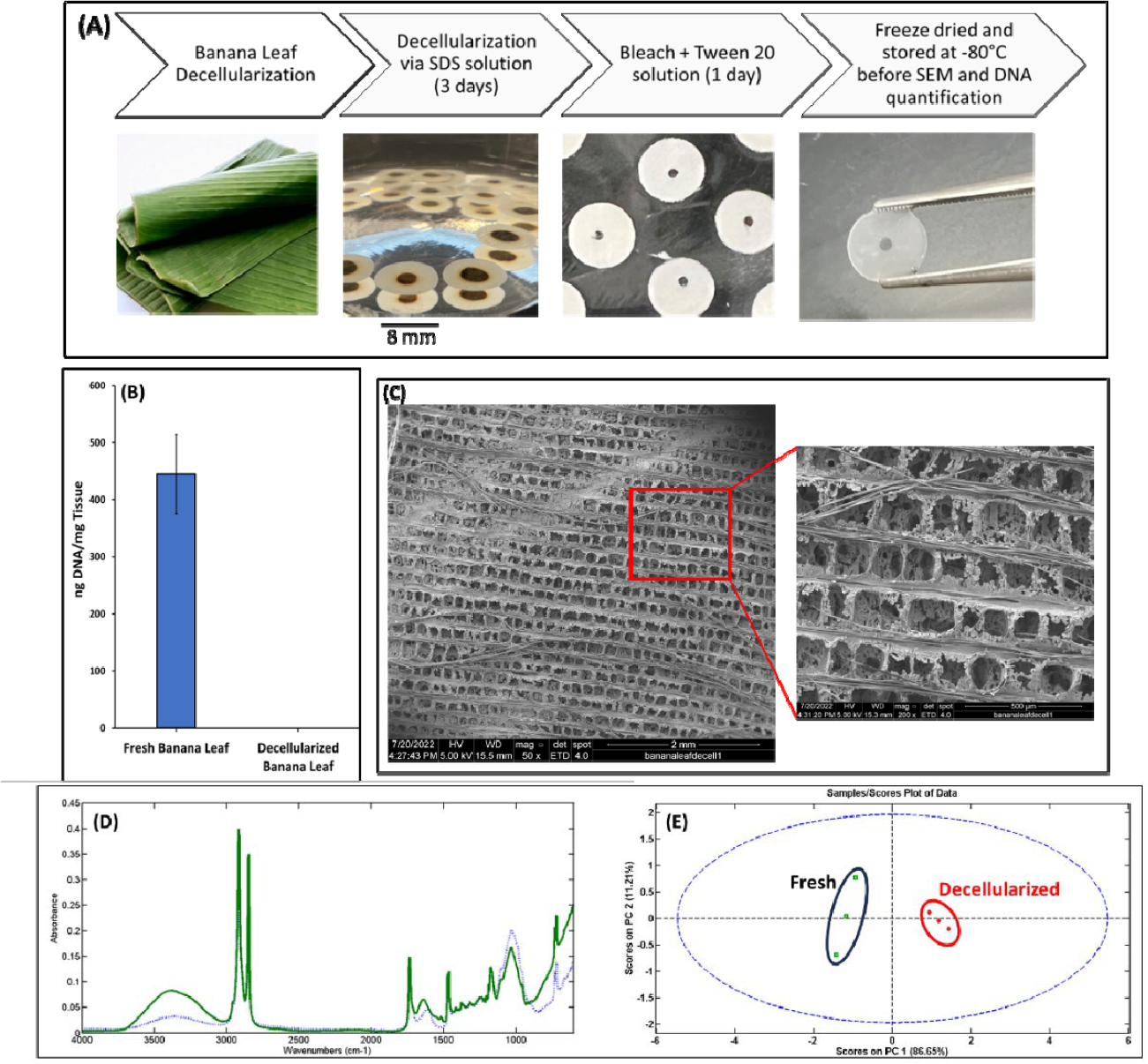
(A) Decellularization process description from fresh banana leaf to completely decellularized translucent banana leaf; (B) Total DNA of fresh and decellularized banana leave (n = 9); (C) SEM pictures of decellularized banana leaves with mean pores size of 147 *μ*m; (D) FTIR spectra of fresh (green) and decellularized (blue) banana leaves; (E) PCA model for fresh and decellularized banana leaves.

The spectra of the banana leaf (green line) and the decellularized banana leaf (blue dots) are illustrated in Figure 1D. The results indicated a decrease in peak intensity and the elimination of certain peaks within the spectral range 1350 cm^-1^ to 1220 cm^-1^ in the decellularized banana leaf, which corresponds to the presence of DNA and RNA peaks. Furthermore, the peak observed at approximately 1520 cm^-1^, commonly associated with cytosine, guanine or amide II, is absent in the decellularized banana leaf. The absence of a peak corresponding to the amide or adenine group was noted at 1650 cm^-1^ in the decellularized banana leaf. Furthermore, a significant decrease in peak intensity was observed at approximately 2950 cm^-1^ in the decellularized banana leaf sample, which primarily corresponds to the stretching of proteins CH bonds (Mello and Vidal, 2012). The results from loading plots also confirmed that the protein and DNA peak intensities accounted for the difference in fresh and decellularized banana leaves.

### Cell Seeding Efficiency

Two distinct methods for cell seeding were employed including natural seeding by directly pipetting the seeding solution onto each scaffold and allowing an incubation phase of 30 minutes, and vacuum infusion. The imaging data suggested that vacuum infusion was more efficient for cell seeding in comparison to conventional seeding (Figure 2). Thus, cell seeding through vacuum perfusion was selected for the rest of the experiments.

**Figure 2.**
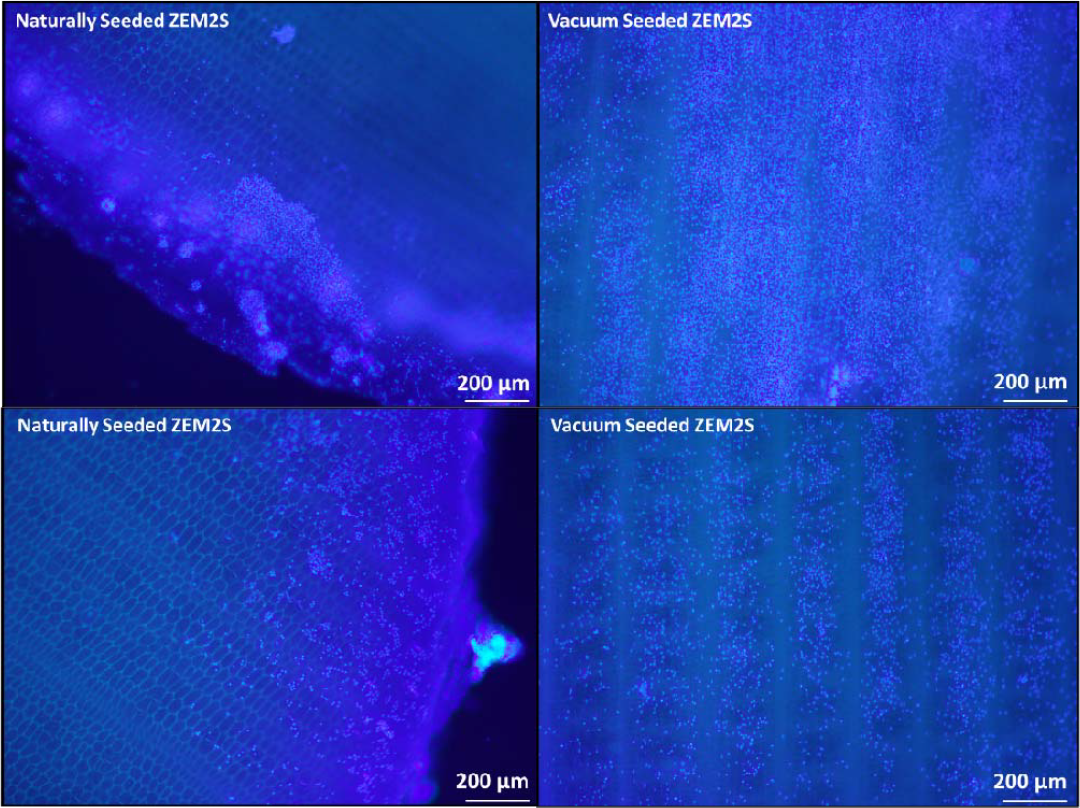
DAPI image from naturally and vacuum seeded cells on the surface of the banana leaves, immediately after seeding. These images indicate high cell biomass around the edges of the decellularized banana leaves, in naturally seeded samples, compared to the uniform cell distribution through the scaffold surface in vacuum seeded cells. Both scaffolds used were uncoated with the thickness of 2000 *μ*m.

The cell seeding efficiency for the vacuum seeding method across all scaffold groups including uncoated, soy-coated, and gelatin-coated (n=27) were 35, 29 and 33% of the total cells in the seeding solution. Following cell seeding, the DNA content in the decellularized banana leave without coatings, as well as those coated with soy and gelatin, were 107, 99, and 88 ng/mg tissue, respectively. These results indicate that the decellularized banana leaf had the capacity to absorb cells up to approximately 25% of its original (pre-decellularization capacity; 445 ng/mg). Cell seeding efficiency was lower than previous findings due to the smaller pore sizes of the matrix compared to cell size. Overall, using vacuum process contributed to the higher DNA content after cell seeding.

### Short-term cell growth indicated by DNA content

DNA quantification indicated no significant difference in cell seeding among all time 0 group (Figure 3). After 24 hrs, the soy coated scaffolds (262.5 ±97 ng/mL) had a significantly higher DNA content than the gelatin coated group (173.63±48.6 ng/mL) (*P < 0*.*05*). DNA quantification indicated that cell growth was reduced between 24 and 48 h. In some previous studies, a reduction in cell numbers after 7 days of incubation was not observed (Jones et al., 2021). Likely factors contributing to these discrepancies may include the method used for cell seeding and the thickness of the scaffold which was around 2,000 *μ*m. It is conceivable that the high cell density and thickness of the decellularized banana leaves used in this study may restrict some nutrient access for the cells, leading to a reduction in cell numbers. However, the technique employed still represents a viable method for effectively seeding cell biomass onto the scaffolds.

**Figure 3.**
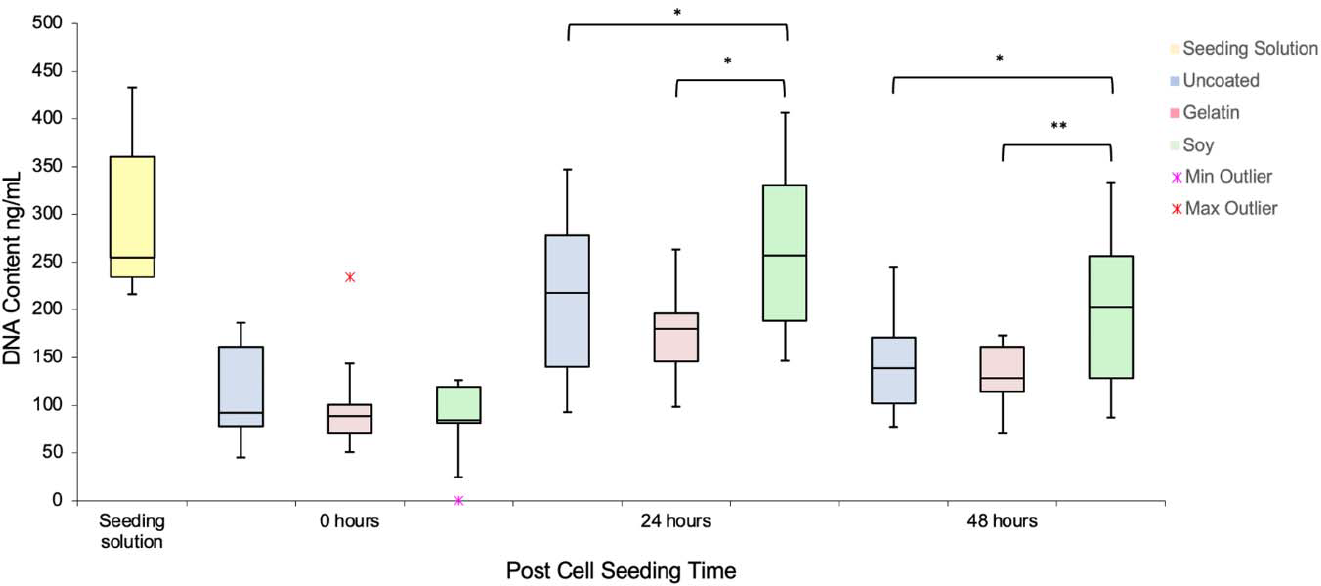
DNA Content over 48 hours in uncoated, soy- and gelatin-coated scaffolds a compared to the initial seeding solution distributed onto each scaffold. Minimum outlier indicates that one of the replicates in soy coated samples had non-detectable level of DNA; Maximum outlier in gelatin-coated samples illustrates that one of the replicates showed 230 ng/mL DNA.

### Cell viability and differentiation efficiency

After 48 hours of incubation in cell growth media, all samples displayed more than 98% viability (Figure 4D). When examining cell adhesion, the soy-coated samples exhibited higher levels of cell adhesion. Conversely, in the case of the gelatin-coated samples, although there was less cell adhesion, the pattern of adhesion appeared more aligned and linear when compared to the soy-coated and uncoated samples (Figure 4, and 5). This observation may be attributed to the fact that gelatin filled all the gaps in the porous decellularized banana leaves, creating grooved areas for the cells to attach. Jones et al. (2021) utilized a gelatin-coated plate as a control group for comparison with decellularized spinach and reported 100% cell viability for both samples. Cell differentiation efficiency using PAX7 indicated that the cells on uncoated and gelatin-coated banana leaves had differentiated more compared to the soy-coated samples. Other researchers also reported delayed differentiation in cells seeded on plant scaffold (Jones et al., 2021). Delayed differentiation potentially suggests increased proliferation and self-renewal. This would lead to an increase in the number of cells and their colonization of a much larger area within the scaffold in the first few days (short term). Additionally, a higher number of the cells would differentiate into myofibers (Riederer et al., 2012).

**Figure 4.**
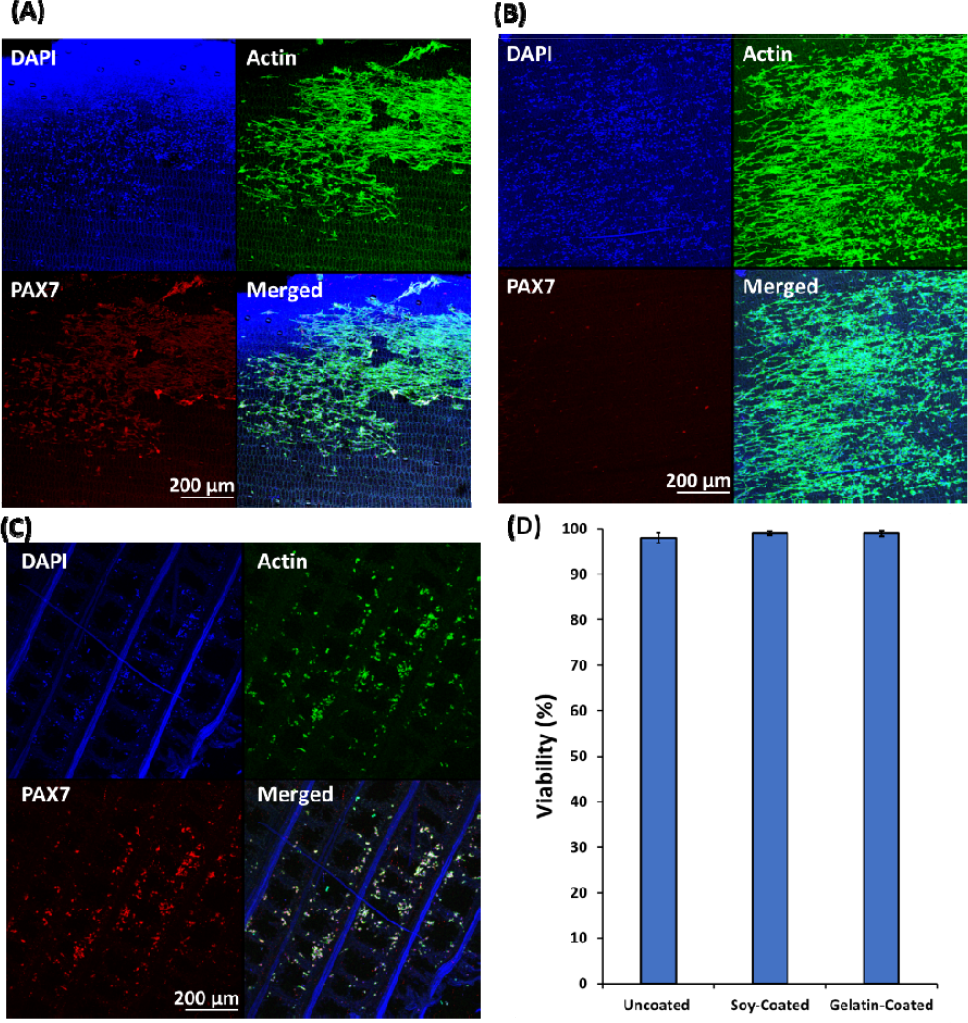
DAPI, Action, PAX7 and merged images for Zebrafish cells vacuum seeded on: (A) uncoated decellularized banana leaves; (B) soy-coated decellularized banana leaves; (C) gelatin-coated decellularized banana leaves; (D) Cell viability determined using Live/Dead kit.

**Figure 5.**
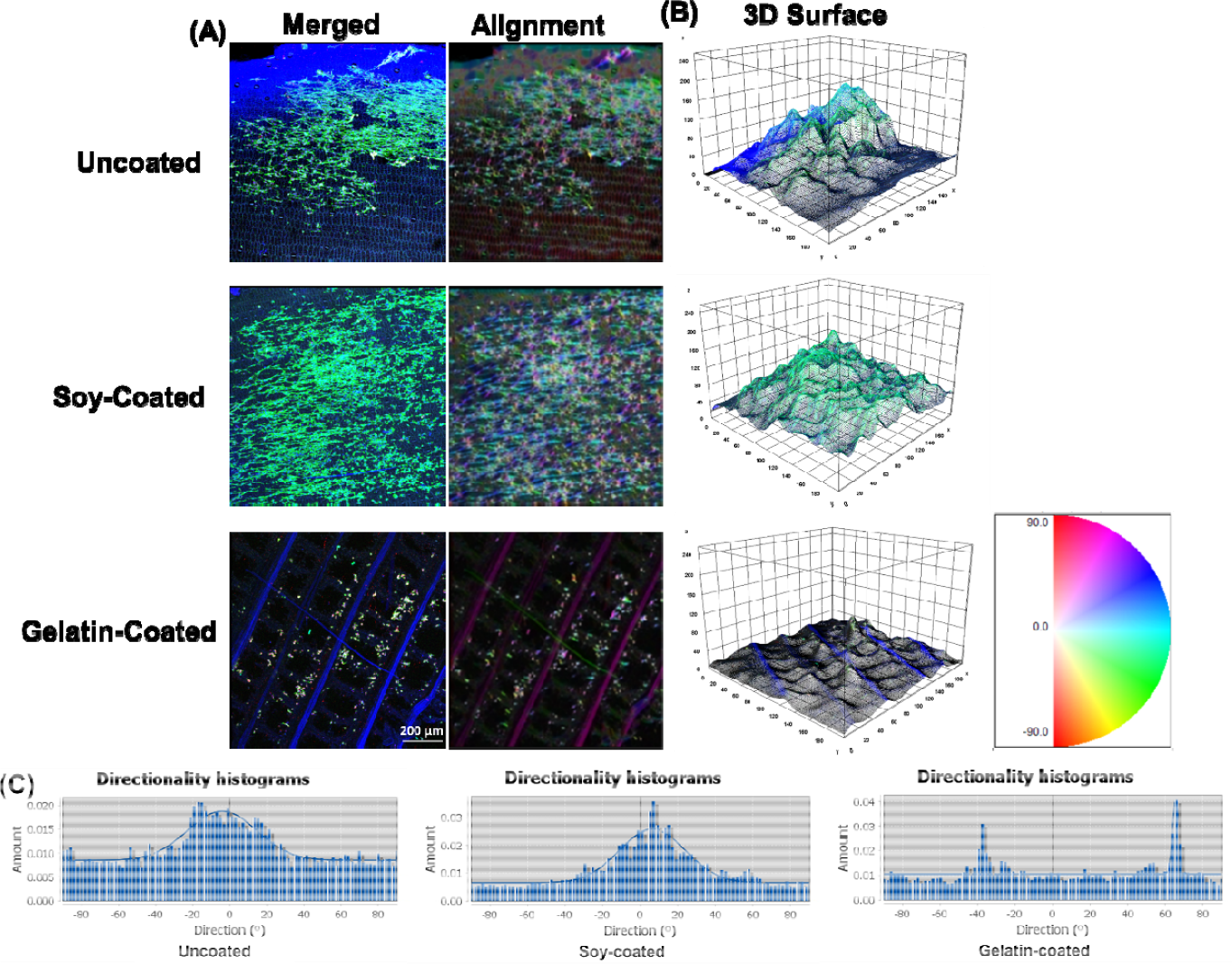
(A) Cell alignment color survey indicating the direction of Zebrafish cytoskeleton on uncoated, soy-coated and gelatin-coated scaffolds; (B) 3D surface analysis of the cytoskeleton filaments on uncoated, soy-coated and gelatin-coated scaffolds; (C) Cytoskeleton microfilament orientation histogram of Zebrafish cells cultured on uncoated, soy-coated, and gelatin-coated banana leaves; (D) The orientation degree ranges from −90 to +90, defining the histograms (C). Any number above 0 indicates that the orientation is positive towards 90, and any degree below 0 is considered as negative towards −90.

The results of alignment indicated that the cells did not align on the un-coated scaffolds (Figure 5), similar to the findings of other researchers (Jones et al., 2021). However, the addition of soy and gelatin as coating materials enhanced cell alignment. Specifically, the dominant orientation based on the image processing results, were −6.3°, 9.2°, and 65.8° for uncoated, soy-coated, and gelatin-coated samples, respectively (Figure 5C).

3D surface plots revealed that the uncoated banana leaves were rougher than the soy- and gelatin-coated samples due to the porous structure. Cell alignment on plant-based materials is influenced by various factors, including surface topography of the decellularized plant tissues, the tissue source and the type of cells (Fontana et al., 2017; Jones et al., 2021). Fontana et al. (2017) highlighted that distinct plant tissues with varying topographical properties can have a significant impact on cell adhesion and alignment. Furthermore, Jones et al. (2021) suggested that plant leaves with grooved topographical features hold promise as potential cultivated meat scaffolds. Recent findings from other researchers have indicated that cells from the same tissue of cows, with a similar age, sex, and breed, may even exhibit different alignment behavior on plant-based scaffolds due to variations in their respective growth environments (Jones et al., 2021). These insights underscore the complexity of cell behavior on plant-based materials and the need for further investigation. This insight would provide valuable insights into optimizing scaffold designs and coating strategies for cultivated seafood production, ultimately advancing the field of cellular agriculture.

### Area shrinkage and activation energy

Area shrinkage is considered one of the most important quality attributes in meat and seafood products in response to heat (Ovissipour et al., 2013, 2017). Area shrinkage is the result of cook loss due to the protein denaturation and aggregation in meat products (Ovissipour et al., 2013, 2017). The results of fresh, cooked and fried scaffolds with the cells are illustrated in Figure 6 A-E. The samples from cooking were used to assess tissue shrinkage (Figure 6 I-IV, Table 2). In plain uncoated and soy-coated decellularized banana leaves, swelling or increase in the area of the samples was observed which was significantly less in soy-coated (Figure 6 III) samples compared to uncoated samples (Figure 6 I). By seeding the cells to the scaffold, different results were observed. The area shrinkage in cell seeded samples was similar to conventional seafood products, in which by increasing the temperature, the area shrinkage was increased (Ovissipour et al., 2013, 2017). Area shrinkage is the result of cook loss due to the protein denaturation and aggregation in meat products (Ovissipour et al., 2013, 2017). After decellularization, the protein content of the scaffold will be significantly reduced, leaving no protein available. This can result in the swelling of carbohydrates in decellularized plants during thermal processing. However, in cell-seeded samples, the protein and moisture contributed significantly to protein denaturation and moisture loss, resulting in high area shrinkage in thermally processed samples. Additionally, the area shrinkage was higher in soy-coated samples due to the availability of more protein for denaturation and the higher cell density on soy-coated samples compared to the uncoated samples.

**Table 2:** Kinetic parameters of area shrinkage of zebrafish cells seeded on uncoated and soy-coated banana leaves.

| Coating status | Kinetics order | Temp (°C) | $K$ (min<sup>-1</sup>) | $R^2$ | $E_a$<br>(kJ/mol) | $K_o$ (min<sup>-1</sup>) | $R^2$ |
| --- | --- | --- | --- | --- | --- | --- | --- |
| Uncoated | Scaffold (First order) | 60 | 0.0118 | 0.87 | 130 ± 9 | 7 × 10<sup>15</sup> | 0.91 |
|  |  | 65 | 0.0347 | 0.80 |  |  |  |
|  |  | 70 | 0.0358 | 0.81 |  |  |  |
|  |  | 75 | 0.0939 | 0.68 |  |  |  |
|  | Cell-Seeded Scaffold<br>(Second order) | 60 | 0.00008 | 0.6 | 195 ± 8 | 4 × 10<sup>23</sup> | 0.90 |
|  |  | 65 | 0.0001 | 0.94 |  |  |  |
|  |  | 70 | 0.0007 | 0.81 |  |  |  |
|  |  | 75 | 0.001 | 0.80 |  |  |  |
| Soy-Coated | Scaffold (First order) | 60 | 0.0308 | 0.60 | 118 ± 11 | 4 × 10<sup>13</sup> | 0.82 |
|  |  | 65 | 0.0366 | 0.72 |  |  |  |
|  |  | 70 | 0.0426 | 0.70 |  |  |  |
|  |  | 75 | 0.2155 | 0.86 |  |  |  |
|  | Cell-Seeded Scaffold<br>(Second order) | 60 | 0.0002 | 0.85 | 198 ± 10 | 7 × 10<sup>23</sup> | 0.73 |
|  |  | 65 | 0.0004 | 0.62 |  |  |  |
|  |  | 70 | 0.0005 | 0.60 |  |  |  |
|  |  | 75 | 0.0053 | 0.82 |  |  |  |

**Figure 6.**
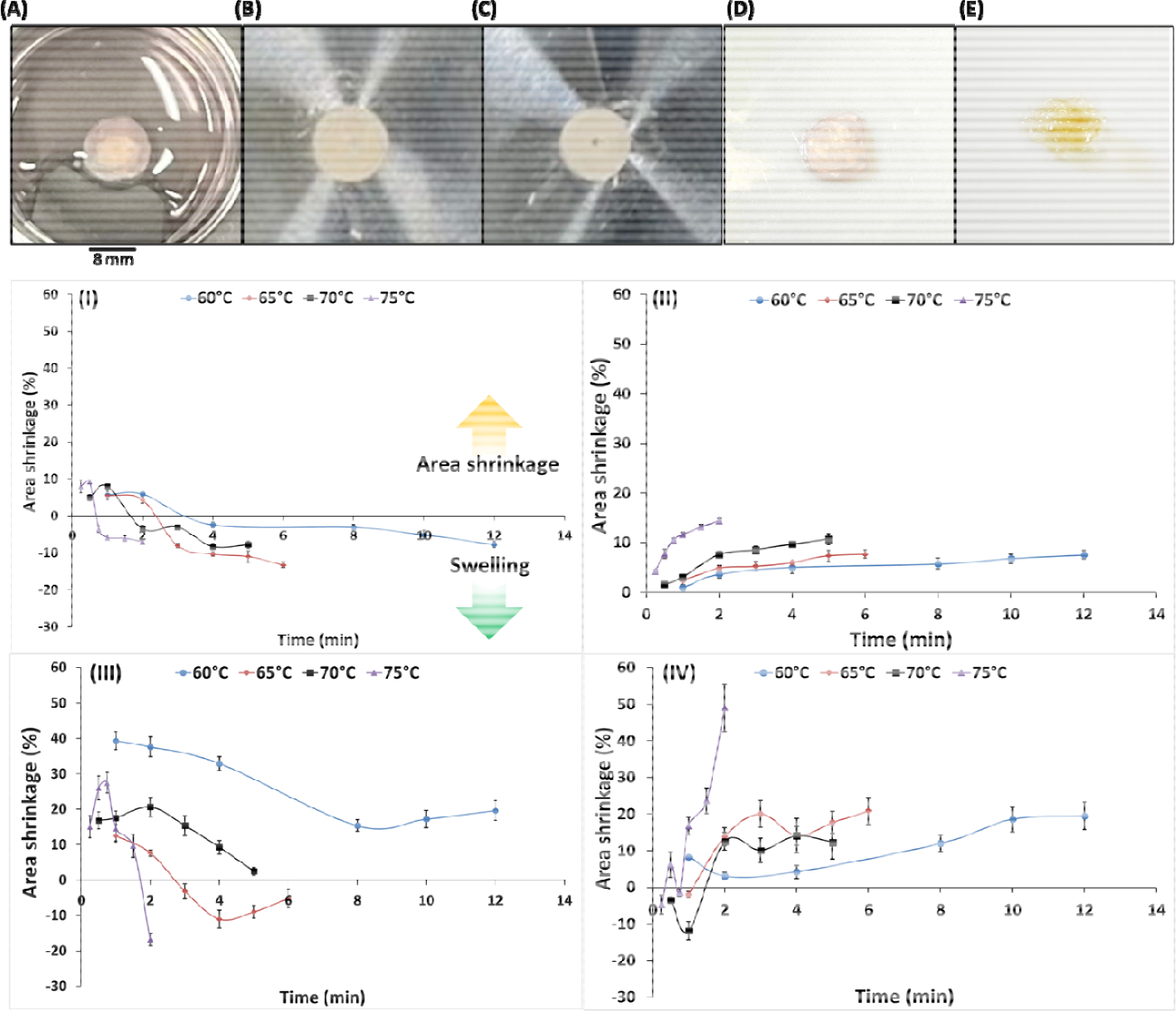
Decellularized banana leaf scaffolds seeded with cells: (A) fresh; (B) before cooking; (C) after cooking at 75°C; (D) fresh sample before frying; (E) after frying; and area shrinkage (%) of (I) uncoated decellularized banana leaf; (II) cell seeded uncoated decellularized banana leaf; (III) soy-coated decellularized banana leaf; (IV) cell seeded soy-coated decellularized banana leaf. The positive areas illustrate the area shrinkage, and the negative areas indicate the swelling and increase in the area due to the impact of heat on the scaffold and possible gelation.

Area shrinkage increased with time and temperature of thermal processing. Area shrinkage in the samples without cells followed the first order with activation energy of 130 and 118 kJ/mol for uncoated, and soy-coated scaffolds, respectively. However, after cell seeding into the scaffolds, the kinetics of area shrinkage shifted to second order, with significantly higher activation energy, reaching 195 and 198 for uncoated and soy-coated scaffolds, respectively. Coating the scaffold with soy protein did not significantly change the activation energy. The results indicated that the activation energy for decellularized banana leaf scaffold, without cell seeding, was relatively high compared to conventional shrinkage area of livestock meat activation energy and was significantly increased after seeding cells on to the scaffolds. Activation energy for area shrinkage in blue mussel (*Mytilus edulis*) (Ovissipour et al., 2013) and Atlantic salmon (*Salmo salar*) (Ovissipour et al., 2017) were 92 and 107 kJ/mol, respectively.

In conventional livestock meat products, heating results in denaturation of myosin and shrinkage of myofibrils, expanding extracellular spaces and resulting in the expulsion of water (Ofstad et al., 1993). In these conventional meat products, muscle fiber diameters and sarcomere length become shorter due to protein denaturation, resulting in water soluble proteins and fats being expelled from the tissue (Bertola et al., 1994; Kong et al., 2007a,b; Ovissipour et al., 2017). In previous studies on area shrinkage in aquatic food products, significant area shrinkage was observed due to thermal processing in salmon (*Oncorhynchus gorbuscha*) (Kong et al., 2007a,b), cod (*Gadus morhua*) (Skipnes et al., 2007, 2011), Atlantic salmon (*S. salar*) (Ofstad et al., 1993; Ovissipour et al., 2017), and blue mussel (*M. edulis*) (Ovissipour et al., 2013). Conventional aquatic food products contain lower collagen contents (Skipnes et al. 2011), resulting in higher susceptibility to heat and higher area shrinkage compared to conventional livestock meat products (Ovissipour et al., 2017). The magnitude of area shrinkage and activation energy vary for different fish. Aquatic organisms with higher muscle fat and collagen content, and stronger connective tissues such as in salmon, exhibit greater heat resistance with higher activating energy (107 kJ/mol) (Ovissipour et al., 2017) compared to tissues with significantly less fat and collagen content and poor connective tissues such as in mussel (92 kJ/mol) (Ovissipour et al., 2013). The findings in the present study present a promising outcome for the use of plant-based scaffolds in cultivating heat-resistant muscle cells, due to the lower area shrinkage with heating. This could potentially lead to the development of cell-based fish fillets that maintain quality, nutritional value, and structural integrity during cooking. The observed high activation energy in decellularized banana leaves further supports the potential of such scaffolds to enhance thermal stability and overall cooking characteristics of cell-based seafood products. Further research and development in this area could impact the cultivated meat industry by providing consumers with sustainable, high-quality alternatives to traditional seafood products.

## Conclusions

Banana leaves offer a cost-effective and sustainable material for scaffold development for cultured meat production. Their potential for decellularized plant tissues for scaffolds in cultivated meat products is an emerging technology. Traditional cell seeding on scaffolds often suffers from inefficiency, necessitating separate bioreactor setups for cell proliferation and differentiation. The present results demonstrate that the use of vacuum improves cell infusion into scaffolds and significantly enhanced cell seeding efficiency, obviates the need for separate bioreactor setups, and has potential to thereby reduce production costs. Furthermore, the kinetics of quality degradation after thermal processing (e.g., cooking) reveal that the activation energy of the final products fall within the range of conventional seafood quality kinetics, suggesting that banana leaves can serve as a scaffold source for cell-based seafood and meat production.

## Acknowledgements

This research was financially supported by the Agriculture and Food Research Initiative (AFRI) Sustainable Agricultural Systems program, grant no. 2021-699012-35978 from the USDA National Institute of Food and Agriculture, and Texas A&M AgriLife Research. We would like to extend our gratitude to Dr. Joseph Awika and Dr. Youjun Deng from Texas A&M University for their invaluable support throughout the course of this study.

## References

Bomkamp, C., Skaalure, S. C., Fernando, G. F., BenLArye, T., Swartz, E. W., & Specht, E. A. (2022). Scaffolding biomaterials for 3D cultivated meat: prospects and challenges. Advanced Science, 9(3), 2102908. 10.1002/advs.202102908

Buizer, A. T., Veldhuizen, A. G., Bulstra, S. K., & Kuijer, R. (2014). Static versus vacuum cell seeding on high and low porosity ceramic scaffolds. Journal of biomaterials applications, 29(1), 3–13. 10.1177/0885328213512171

Cheng, Y.-W., Shiwarski, D. J., Ball, R. L., Whitehead, K. A., & Feinberg, A. W. (2020). Engineering aligned skeletal muscle tissue using decellularized plant-derived scaffolds. ACS Biomaterials Science & Engineering, 6(5), 3046–3054. 10.1021/acsbiomaterials.0c00058

Crapo, P. M., Gilbert, T. W., & Badylak, S. F. (2011). An overview of tissue and whole organ decellularization processes. Biomaterials, 32(12), 3233–3243. 10.1016/j.biomaterials.2011.01.057

Eibl, R., Senn, Y., Gubser, G., Jossen, V., van den Bos, C., & Eibl, D. (2021). Cellular agriculture: Opportunities and challenges. Annual Review of Food Science and Technology, 12, 51–73. 10.1146/annurev-food-063020-123940

Gershlak, J. R., Hernandez, S., Fontana, G., Perreault, L. R., Hansen, K. J., Larson, S. A., Binder, B. Y., Dolivo, D. M., Yang, T., & Dominko, T. (2017). Crossing kingdoms: Using decellularized plants as perfusable tissue engineering scaffolds. Biomaterials, 125, 13–22. 10.1016/j.biomaterials.2017.02.011

Jones, J. D., Rebello, A. S., & Gaudette, G. R. (2021). Decellularized spinach: An edible scaffold for laboratory-grown meat. Food Bioscience, 41, 100986. 10.1016/j.fbio.2021.100986

Jones, J. D., Thyden, R., Perreault, L. R., Varieur, B. M., Patmanidis, A. A., Daley, L., Gaudette, G. R., & Dominko, T. (2023). Decellularization: Leveraging a Tissue Engineering Technique for Food Production. ACS Biomaterials Science & Engineering, 9(5), 2292–2300. 10.1021/acsbiomaterials.2c01421

Kong, F., Tang, J., Rasco, B., & Crapo, C. (2007). Kinetics of salmon quality changes during thermal processing. Journal of food engineering, 83(4), 510–520. 10.1016/j.jfoodeng.2007.04.002

Kong, F., Tang, J., Rasco, B., Crapo, C., & Smiley, S. (2007). Quality changes of salmon (Oncorhynchus gorbuscha) muscle during thermal processing. Journal of food science, 72(2), S103–S111. 10.1111/j.1750-3841.2006.00246.x

Kumar, P., Sharma, N., Sharma, S., Mehta, N., Verma, A. K., Chemmalar, S., & Sazili, A. Q. (2021). In-vitro meat: a promising solution for sustainability of meat sector. Journal of animal science and technology, 63(4), 693. 10.5187%2Fjast2021.e85

Lanzoni, D., Bracco, F., Cheli, F., Colosimo, B. M., Moscatelli, D., Baldi, A., Rebucci, R., & Giromini, C. (2022). Biotechnological and technical challenges related to cultured meat production. Applied Sciences, 12(13), 6771. 10.3390/app12136771

Mello, M. L. S., & Vidal, B. (2012). Changes in the infrared microspectroscopic characteristics of DNA caused by cationic elements, different base richness and single-stranded form. PLoS One. 10.1371/journal.pone.0043169

Nichol, J. W., & Khademhosseini, A. (2009). Modular tissue engineering: engineering biological tissues from the bottom up. Soft matter, 5(7), 1312–1319. 10.1039/b814285h

Ofstad, R., Kidman, S., Myklebust, R., & Hermansson, A.-M. (1993). Liquid holding capacity and structural changes during heating of fish muscle: cod (Gadus morhua L.) and salmon (Salmo salar). Food structure, 12(2), 4.

Ovissipour, M., Rasco, B., Tang, J., & Sablani, S. S. (2013). Kinetics of quality changes in whole blue mussel (Mytilus edulis) during pasteurization. Food Research International, 53(1), 141–148. 10.1016/j.foodres.2013.04.029

Ovissipour, M., Rasco, B., Tang, J., & Sablani, S. (2017). Kinetics of protein degradation and physical changes in thermally processed Atlantic salmon (Salmo salar). Food and bioprocess technology, 10, 1865–1882. 10.1007/s11947-017-1958-4

Perreault, L. R., Thyden, R., Kloster, J., Jones, J. D., Nunes, J., Patmanidis, A. A., Reddig, D., Dominko, T., & Gaudette, G. R. (2023). Repurposing agricultural waste as low-cost cultured meat scaffolds. Frontiers in Food Science and Technology, 3, 1208298. 10.3389/frfst.2023.1208298

Püspöki, Z., Storath, M., Sage, D., & Unser, M. (2016). Transforms and operators for directional bioimage analysis: a survey. Focus on bio-image informatics, 69–93. 10.1007/978-3-319-28549-8_3

Rezakhaniha, R., Agianniotis, A., Schrauwen, J. T. C., Griffa, A., Sage, D., Bouten, C. v., Van De Vosse, F., Unser, M., & Stergiopulos, N. (2012). Experimental investigation of collagen waviness and orientation in the arterial adventitia using confocal laser scanning microscopy. Biomechanics and modeling in mechanobiology, 11, 461–473.10.1007/s10237-011-0325-z

Riederer, I., Negroni, E., Bencze, M., Wolff, A., Aamiri, A., Di Santo, J. P., Silva-Barbosa, S. D., Butler-Browne, G., Savino, W., & Mouly, V. (2012). Slowing down differentiation of engrafted human myoblasts into immunodeficient mice correlates with increased proliferation and migration. Molecular Therapy, 20(1), 146–154. 10.1038/mt.2011.193

Rischer, H., Szilvay, G. R., & Oksman-Caldentey, K.-M. (2020). Cellular agriculture—industrial biotechnology for food and materials. Current opinion in biotechnology, 61, 128–134. 10.1016/j.copbio.2019.12.003

Röös, E., Bajželj, B., Smith, P., Patel, M., Little, D., & Garnett, T. (2017). Greedy or needy? Land use and climate impacts of food in 2050 under different livestock futures. Global Environmental Change, 47, 1–12. 10.1016/j.gloenvcha.2017.09.001

Schindelin, J., Arganda-Carreras, I., Frise, E., Kaynig, V., Longair, M., Pietzsch, T., Preibisch, S., Rueden, C., Saalfeld, S., & Schmid, B. (2012). Fiji: an open-source platform for biological-image analysis. Nature methods, 9(7), 676–682. 10.1038/nmeth.2019

Skipnes, D., Østby, M. L., & Hendrickx, M. E. (2007). A method for characterising cook loss and water holding capacity in heat treated cod (Gadus morhua) muscle. Journal of food engineering, 80(4), 1078–1085. 10.1016/j.jfoodeng.2006.08.015

Skipnes, D., Johnsen, S., Skåra, T., Sivertsvik, M., & Lekang, O. (2011). Optimization of heat processing of farmed Atlantic cod (Gadus morhua) muscle with respect to cook loss, water holding capacity, color, and texture. Journal of Aquatic Food Product Technology, 20(3), 331–340. 10.1080/10498850.2011.571808

Soice, E., & Johnston, J. (2021). How cellular agriculture systems can promote food security. Frontiers in Sustainable Food Systems, 5, 753996. 10.3389/fsufs.2021.753996

Stephens, N., Di Silvio, L., Dunsford, I., Ellis, M., Glencross, A., & Sexton, A. (2018). Bringing cultured meat to market: Technical, socio-political, and regulatory challenges in cellular agriculture. Trends in Food Science & Technology, 78, 155–166. 10.1016/j.tifs.2018.04.010

Thyden, R., Perreault, L. R., Jones, J. D., Notman, H., Varieur, B. M., Patmanidis, A. A., Dominko, T., & Gaudette, G. R. (2022). An edible, decellularized plant derived cell carrier for lab grown meat. Applied Sciences, 12(10), 5155. 10.3390/app12105155

